# Analysis of the mutation dynamics of SARS-CoV-2 reveals the spread history and emergence of RBD mutant with lower ACE2 binding affinity

**DOI:** 10.1101/2020.04.09.034942

**Authors:** Yong Jia, Gangxu Shen, Stephanie Nguyen, Yujuan Zhang, Keng-Shiang Huang, Hsing-Ying Ho, Wei-Shio Hor, Chih-Hui Yang, John B Bruning, Chengdao Li, Wei-Lung Wang

## Abstract

Monitoring the mutation dynamics of SARS-CoV-2 is critical for the development of effective approaches to contain the pathogen. By analyzing 106 SARS-CoV-2 and 39 SARS genome sequences, we provided direct genetic evidence that SARS-CoV-2 has a much lower mutation rate than SARS. Minimum Evolution phylogeny analysis revealed the putative original status of SARS-CoV-2 and the early-stage spread history. The discrepant phylogenies for the spike protein and its receptor binding domain proved a previously reported structural rearrangement prior to the emergence of SARS-CoV-2. Despite that we found the spike glycoprotein of SARS-CoV-2 is particularly more conserved, we identified a receptor binding domain mutation that leads to weaker ACE2 binding capability based on in silico simulation, which concerns a SARS-CoV-2 sample collected on 27^th^ January 2020 from India. This represents the first report of a significant SARS-CoV-2 mutant, and requires attention from researchers working on vaccine development around the world.

**Highlights:**

- Based on the currently available genome sequence data, we provided direct genetic evidence that the SARS-COV-2 genome has a much lower mutation rate and genetic diversity than SARS during the 2002-2003 outbreak.
- The spike (S) protein encoding gene of SARS-COV-2 is found relatively more conserved than other protein-encoding genes, which is a good indication for the ongoing antiviral drug and vaccine development.
- Minimum Evolution phylogeny analysis revealed the putative original status of SARS-CoV-2 and the early-stage spread history.
- We confirmed a previously reported rearrangement in the S protein arrangement of SARS-COV-2, and propose that this rearrangement should have occurred between human SARS-CoV and a bat SARS-CoV, at a time point much earlier before SARS-COV-2 transmission to human.
- We provided first evidence that a mutated SARS-COV-2 with reduced human ACE2 receptor binding affinity have emerged in India based on a sample collected on 27th January 2020.

## Introduction

The outbreak of severe acute respiratory syndrome–coronavirus 2 (SARS-CoV-2) has caused an unprecedented pandemic and severe fatality around the world. As of 4^th^ April 2020, the total number of SARS-CoV-2 infection has reached over 1.05 million cases globally, claiming 56,985 lives (Coronavirus disease 2019, Situation Report-15, WHO). Most concerning is that this number is predicted to continue to rise significantly in the coming months. Scientists have been working diligently to understand how the virus spreads and evolves. There is an imminent challenge to develop effective approaches to contain the rapid spread of this pathogen.

In addition to the traditional control methods, such as travel bans and self-isolation, which have clear negative impacts on the economy and disrupt normal social activities, the development of antiviral drugs and vaccines should be the ultimate solution to contain the epidemic and reduce the fatality^1,2^. Similar to other SARS-like CoVs ^3,4^, SARS-CoV-2 uses its spike (S) protein to bind and invade human cells ^5,6^. The S protein and its host receptor are the key targets for drug design and vaccine development ^7,8^. Recently, several 3D protein structures of the receptor binding domain (RBD) of SARS-CoV-2 spike protein have been determined ^5,6,9^. Elucidation of the structural basis of receptor recognition by SARS-CoV-2 has laid the foundation for future vaccine development ^6,9^.

Vaccines utilize the human immune system and is specific to the viral-encoded peptides ^10^. One of the major concerns for antiviral vaccine development is the constant emergence of new mutations, which may reduce its efficacy in future epidemics ^7,10^. A prominent example is Influenza virus in which mutations arise every year, requiring annual immunization ^11^. SARS-CoV-2 is a single-stranded RNA virus, whose genome can readily mutate as the virus spreads ^12,13^. Interestingly, initial assessment of the first 9 SARS-CoV-2 genome sequences revealed a low level of mutation rate ^14^. Several more recent studies also highlighted relatively low genetic diversity and stable genomes for SARS-CoV-2 ^15–17^, which suggests that only a single vaccine may be required for SARS-CoV-2. However, these results may be based on limited genomic data in the early stage of virus development. It is critical to study and monitor the mutation dynamics of SARS-COV-2 to gain a more accurate understanding of the virus and therefore guide vaccine development.

Taking advantage of the increasing amount of genomic data collected around the world, we set to explore the current status of SARS-CoV-2 genomic diversity, assess the mutation rate, and potentially identify the emergence of novel mutations that may require close attention. A total of 106 complete or near complete SARS-CoV-2 genome data covering over 12 countries was downloaded from a public database. The genetic diversity profile and evolutionary rate for each protein-encoding gene was characterized. Phylogenetic analyses in this study revealed clues to the spread history of SARS-CoV-2 in some countries. Most importantly, we identified a SARS-CoV-2 mutation with likely reduced human angiotensin-converting enzyme 2 (ACE2) binding affinity. We confirmed that SARS-CoV-2 has a relatively low mutation rate and suggest that novel mutation with likely varied virulence and different immune characteristics may also emerge.

## Methods

### Sequence retrieval

The latest sequence data for SARS-CoV-2 and SARS was retrieved from the NCBI public database at https://www.ncbi.nlm.nih.gov/genbank/sars-cov-2-seqs/. The 5’UTR, 3’UTR, and CDS sequences of the reference SARS-CoV-2 genome (NC_045512.2) and the human SARS genome (NC_004718.3) were used to blastn against the available genome data. The homology search targets were restricted to the complete or near-complete genomes for further analyses.

### Conservation profiling

The assessment of sequence conservation was performed using the PLOTCON tool from the The European Molecular Biology Open Software Suite at https://www.bioinformatics.nl/cgi-bin/emboss/plotcon. The gene model of SARS-CoV-2 was generated using the AnnotationSketch ^18^ tool based on the genome annotation data downloaded from NCBI database.

### Phylogeny construction

Codon-based sequence alignment was performed for the conserved domain sequences (CDS) using the MUSCLE program (limited to 2 iterations for fast alignment of long sequences) ^19^. Alignment of the 5’UTR and 3’UTR sequences were performed separately. The obtained alignment files were concatenated for final phylogeny construction. The phylogenetic tree was developed in MEGA7.0 ^20^ using the Minimum Evolution method with p-distance substitution model, and the Maximum Likelihood method (HKY+G+I, 500 times bootstrap test) for the S protein analyses. Tree annotation was carried out using Figtree software (http://tree.bio.ed.ac.uk/software/figtree/) and cophyloplot from ape 5.0 R package ^21^.

### Evolutionary rate assessment

The ratio of nonsynonymous mutations (*d_N_*) to synonymous mutations (*d_S_*) was calculated using codeml in the PAML (version 4.7) package ^22^. CDS sequences for each protein encoding gene were filtered to remove redundant identical sequences. Then codon-based CDS sequence alignment was performed using the MUSCLE program, and an individual NJ tree was generated using MEGA7.0 ^20^ with p-distance model. The obtained sequence alignment and phylogenetic tree files were used as PAML inputs for *d_N_* and *d_S_* calculations.

### Protein structural analyses

3D structure of the SARS-CoV-2 spike glycoprotein in complex with human ACE2 (PDB: 6VW1) has been determined recently ^5,9^. The structural model for the receptor binding domain (RBD) was extracted from 6VW1 for comparison analysis with the human SARS structure (PDB: 2AJF) ^3^, which is in complex with the receptor: human ACE2. Amino acid sequence alignment of the spike glycoprotein was visualized and annotated using ESPript 3.0 tool (http://espript.ibcp.fr/ESPript/ESPript/index.php). Protein hydrophobicity profiles were implemented in PyMOL using the Color_h script (http://www.pymolwiki.org/index.php/Color_h), based on the hydrophobicity scale defined at http://us.expasy.org/tools/pscale/Hphob.Eisenberg.html. All structure visualization was carried out using PyMol (Version 2.2.3. Schrodinger, LLC).

### In silico mutagenesis and prediction of change in binding free energy

The crystal structure of SARS-CoV-2 spike glycoprotein in complex with the human receptor, angiotensin-converting enzyme 2 (ACE2) (PDB: 6VW1) was used to generate a model of the R408I SARS-CoV-2 spike glycoprotein mutant using ICM-Pro (Molsoft LLC, La Jolla, CA, USA). The model was subsequently refined through the optimization of geometric restraints, refinement of clashing side-chains and minimization of free energy. Prediction of the binding free energy change of the SARS-CoV-2 spike glycoprotein, wild-type and R408I mutant, and the ACE2 receptor interaction was performed using ICM-Pro. All structure visualization was carried out using PyMol (Version 2.2.3. Schrodinger, LLC).

## Results

### Genetic diversity analyses identified a single amino acid mutation in RBD of the spike protein in SARS-CoV-2

As of 24^th^ March 2020, there are a total of 174 nucleotide sequences for SARS-CoV-2 in the NCBI database. By restricting to the complete or near-complete genomes, 106 sequences from 12 countries were obtained and used for further analyses. This encompasses 54 records from USA, 35 from China, and the rest from other countries: Australia (1), Brazil (2), Finland (1), India (2), Italy (1), Japan (3), Nepal (1), Spain (3), South Korea (1), and Sweden (1).

Based on the gene model of the reference SARS-CoV-2 genome (GeneBank: NC_045512.2), a total of 12 protein-encoding open reading frames (ORFs), plus the 5’UTR and 3’UTR were annotated (**Figure 1A**). Overall, the gene sequences from different samples are highly homologous, sharing > 99.1% identity, apart from the 5’UTR (96.7%) and 3’UTR (98%) (**Table 1**), which are relatively more divergent. Sequence alignment showed that there is no mutation in ORF6, ORF7a, and ORF7b. The genetic diversity profile across the 106 genomes was displayed in **Figure 1A**. A few nucleotide sites within ORF1a, ORF1b, ORF3a, and ORF8 exhibiting high genetic diversity were identified (**Figure 1A**).

**Table 1.** List of SARS-CoV-2 strains containing the R408I mutation. GISAID and NCBI databases (as of 5^th^ Oct 2020) were searched for strains containing the R408I mutation.

| No | Sample name | Accession ID | Collection date | Location |
| --- | --- | --- | --- | --- |
| 1 | hCoV-19/Switzerland/BE-ETHZ-280249/2020 | EPI_ISL_560454 | 10/09/2020 | Switzerland / Bern |
| 2 | hCoV-19/England/MILK-9AA096/2020 | EPI_ISL_550882 | 1/09/2020 | England |
| 3 | hCoV-19/England/OXON-B1C55/2020 | EPI_ISL_479081 | 30/06/2020 (Submission) | England |
| 4 | hCoV-19/Egypt/CUNCI-HGC3I013/2020<br>(MT627395 NCBI database) | EPI_ISL_479694<br>(MT627395.1) | 2/06/2020 | Egypt |
| 5 | hCoV-19/England/NORT-29DB8D/2020 | EPI_ISL_484309 | 7/05/2020 | England |
| 6 | hCoV-19/England/NORT-29E005/2020 | EPI_ISL_488118 | 6/05/2020 | England |
| 7 | hCoV-19/England/NORW-E7A01/2020 | EPI_ISL_449088 | 4/05/2020 | England |
| 8 | hCoV-19/England/NORT-29D437/2020 | EPI_ISL_499803 | 3/05/2020 | England |
| 9 | hCoV-19/Egypt/CUNCI-HGC013/2020<br>(MT510693 NCBI database) | EPI_ISL_468047<br>(MT510693.1) | 2/05/2020 | Egypt |
| 10 | hCoV-19/England/NORW-EE30F/2020 | EPI_ISL_490529 | 24/04/2020 | England |
| 11 | hCoV-19/England/NORT-29C84B/2020 | EPI_ISL_488132 | 23/04/2020 | England |
| 12 | hCoV-19/England/PORT-2D11E5/2020 | EPI_ISL_475338 | 21/04/2020 | England |
| 13 | hCoV-19/England/OXON-B07DD/2020 | EPI_ISL_478909 | 20/04/2020 | England |
| 14 | hCoV-19/Portugal/PT0716/2020 | EPI_ISL_510975 | 24/03/2020 | Portugal |
| 15 | hCoV-19/England/CAMB-74A09/2020 | EPI_ISL_425342 | 18/03/2020 | England |
| 16 | MT050491 (NCBI database) | MT050491.1 | 30/1/2020 | India / Kerala |
| 17 | hCoV-19/India/MH-1-27/2020<br>(MT012098, NCBI database) | EPI_ISL_413522<br>(MT012098.1) | 27/01/2020 | India / Kerala |

**Figure 1.**
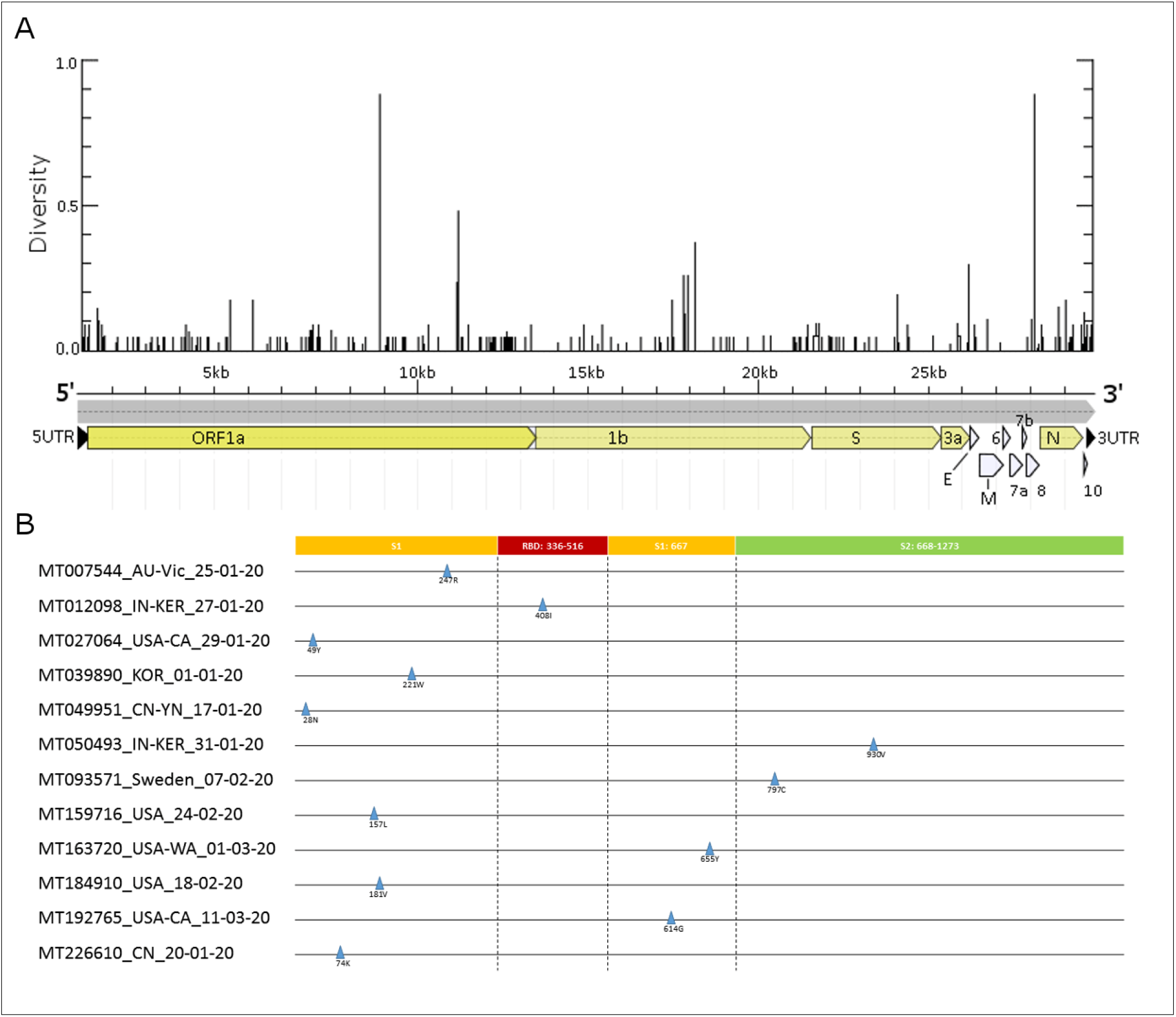
Genetic diversity profile of SARS-CoV-2 genomes and amino acid mutations in the spike glycoprotein. **A**) Pair-wise genetic distance for each nucleotide site calculated from the 106 SARS-CoV-2 genomes. Gene model is based on the reference genome (GeneBank: NC_045512.2). **B**) Identification of amino acid mutations in the spike glycoprotein. Sequences were named as: Accession name_country_ sample collection time (AU: Australia; IN: India; USA: United States; KOR: South Korea; CN: China; Sweden: Sweden.) Amino acid numbering according to the reference spike protein (Accession ID: YP_009724390.1).

The S protein is critical for virus infection and vaccine development. As shown in **Figure 1B**, 12 single amino acid substitutions in 12 genomes were identified for the spike glycoprotein, only one (R408I) of which occurs in the receptor binding domain (RBD). This mutation concerns an accession collected from Kerala State, India on 27^th^ Jan 2020.

To track the occurrence of the R408I mutant, we checked the latest GISAID database (5^th^ Oct 2020) and confirmed that there are a total of 17 SARS-CoV-2 strains containing the R408I mutation (**Table 1**): England (11), Egypt (2), Portugal (1), Switzerland (1), and India (2). We believe that these numbers are still underestimated by the limited sequencing capacity in respective countries. For example, there are only a total of 152 spike protein records for Egypt in the GISAID database. Noteworthy, the latest R408I SARS-CoV-2 samples were collected on 10^th^ Sep 2020 and 1^st^ Sep 2020 from Switzerland and England (**Table 1**), respectively, indicating that this mutant is still actively spreading.

### SARS-CoV-2 displayed a much lower mutation rate than SARS-CoV, with a highly conserved S gene

To assess the mutation rate and genetic diversity of SARS-CoV-2, the ratio of nonsynonymous mutations (*d_N_*) and synonymous mutations (*d_S_*) was calculated for each protein-encoding ORF based on the 106 SARS-CoV-2 and 39 SARS genomes. For SARS-CoV-2, the highest *d_N_* was observed for ORF8 (0.0111), followed by ORF1a (0.0081), ORF9 (0.0079), and ORF4 (0.0063) (**Table 2**), indicating these genes may be more likely to accumulate nonsynonymous mutations. In contrast, ORF1b (0.0029), S gene (0.0040) encoding the spike protein, and ORF5 (0.0023) are relatively more conserved in terms of nonsynonymous mutation. Noteworthy, ORF6, ORF7ab and ORF10 are strictly conserved with no nonsynonymous mutation. Compared to SARS-CoV-2, SARS displayed higher mutation rates for all of the ORFs in the virus genome (Table 1), suggesting an overall higher level of genetic diversity and mutation rate. In particular, the *d_N_* and *d_S_* values for the S gene in SARS-CoV is approximately 12 and 7 times higher than that for SARS-CoV-2, respectively. In contrast, the mutation rate differences for ORF1a and ORF1b between SARS-CoV-2 and SARS are relatively milder, varying from 1.5 times to 4.8 times only. In contrast to SARS-CoV-2, which has strictly conserved ORF6, ORF7a, and ORF7b, SARS displayed mutation rates at different levels. Notably, the *d_S_* for ORF10 are comparable between the two genomes at 0.0326 and 0.0341, respectively.

**Table 2.** Mutation rate analysis on SARS-CoV-2 genes. Gene model is according to the SARS-CoV-2 reference genome (GeneBank: NC_045512.2). S: spike glycoprotein. “Pair-wise identity” indicate the minimum pair-wise sequence identity among the 106 genomes. ****dN****: nonsynonymous mutation; ****dS****: synonymous mutations. “--”: not applicable.

| Gene name | 5'UTR | 1a | 1b | S | ORF3a | ORF4_E | ORF5_M | ORF6 | ORF7a | ORF7b | ORF8 | ORF9 | ORF10 | 3'UTR |
| --- | --- | --- | --- | --- | --- | --- | --- | --- | --- | --- | --- | --- | --- | --- |
| Length (bp) | 211 | 13218 | 8088 | 3822 | 828 | 228 | 669 | 186 | 336 | 132 | 366 | 1260 | 117 | 152 |
| Pair-wise identity | 96.7% | 99.8% | 99.9% | 99.9% | 99.6% | 99.1% | 99.7% | 100% | 100% | 100% | 99.5% | 99.7% | 99.1% | 98% |
| *d<sub>N</sub>* SARS-CoV-2 | -- | 0.0081 | 0.0029 | 0.0040 | 0.0074 | 0.0063 | 0.0023 | 0 | 0 | 0 | 0.0111 | 0.0079 | 0 | -- |
| SARS | -- | 0.0119 | 0.0077 | 0.0532 | 0.0331 | 0.0338 | 0.023 | 0.3031 | 0.0040 | 0.5339 | 0.0287 | 0.0197 | 0.0135 | -- |
| *d<sub>S</sub>* SARS-CoV-2 | -- | 0.0041 | 0.0083 | 0.0055 | 0 | 0.0611 | 0.0046 | 0 | 0 | 0 | 0 | 0.0172 | 0.0326 | -- |
| SARS | -- | 0.0196 | 0.0326 | 0.0442 | 0.0248 | 0.0146 | 0.0928 | 0.0202 | 0.0183 | 0.0005 | 0.0566 | 0.9552 | 0.0341 | -- |

### Phylogeny analysis revealed the original status of SARS-CoV-2 and its spread history

To trace the potential spread history of SARS-CoV-2 across the world, an unrooted Minimum Evolution (ME) tree of the 106 genomes was developed based on whole-genome sequence alignment. The clustering pattern of the ME phylogeny (**Figure 2**) shed light on how the virus may have spread at the early stage. At the center of the ME tree, a number of virus accessions collected from China (including the reference genome NC_045512.2) and USA have the shortest branch (marked by red and black dots), thus may indicate the original status of SARS-CoV-2. The radial pattern, instead of clustering together, of these accessions and other accessions derived from the tree center (highlighted in yellow color) with longer branches, implies the independent mutations occurring during the virus spread (**Figure 2**). However, the longer branch may not be always associated with a longer evolution time, as some accessions collected in December 2019 have equal or even longer branch than those collected in January and February 2020.

**Figure 2.**
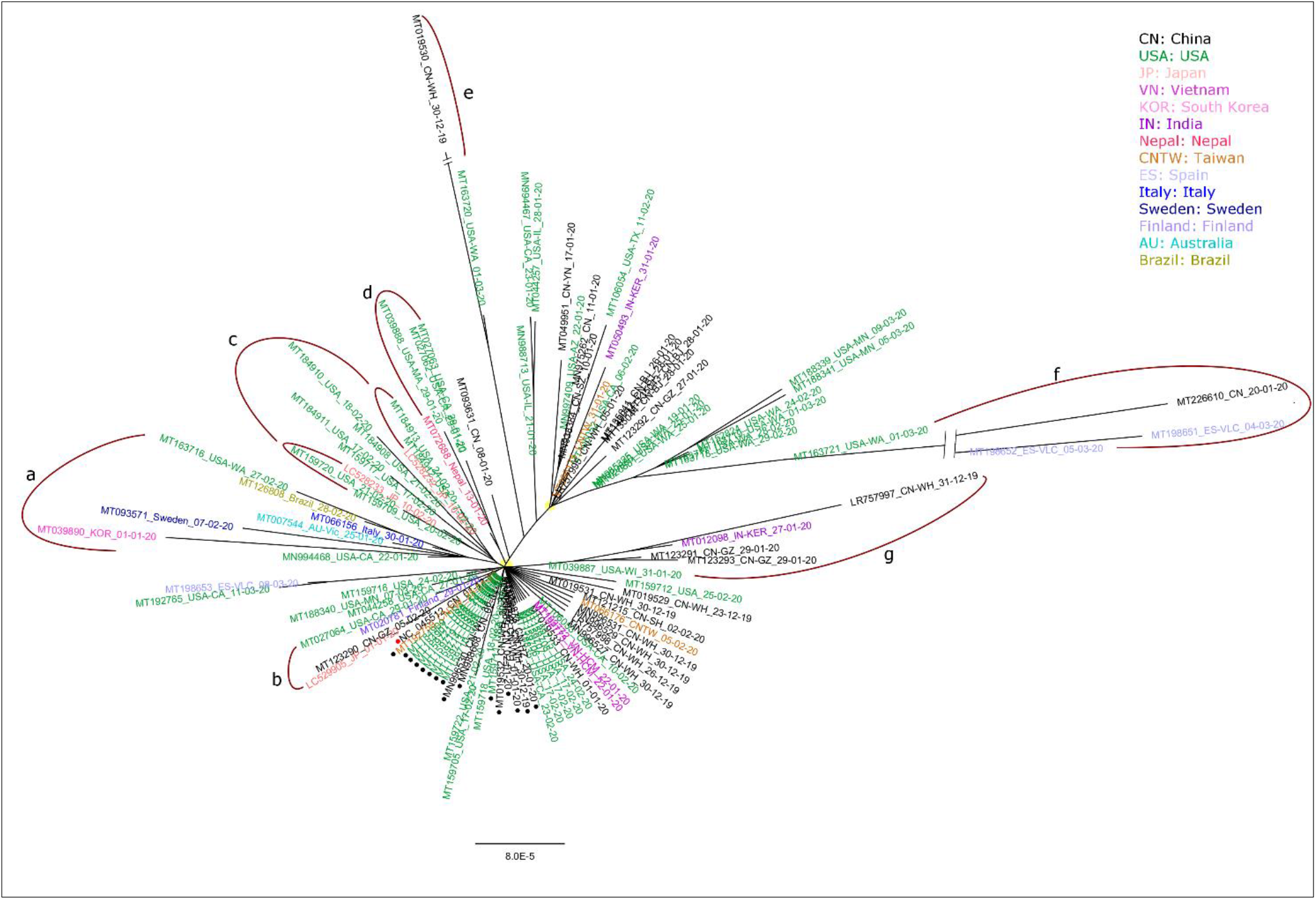
Phylogeny clustering analyses of the 106 SARS-CoV-2 genomes. Results were based on whole genome sequence alignment using Minimum Evolution method. Each accession was named in the “accession ID, country, sample collection time” format. Samples collected from different countries were highlighted in different colors. Red dots indicated the reference SARS-CoV-2 genome (GeneBank: NC_045512.2), which together with black dots indicated the original status of SARS-CoV-2 (branch length = 0). The putative two types of SARS-CoV-2 were highlighted in yellow shades. Notable clades containing sequences from more than one country were highlighted in curved line (magenta).

Due to the data availability, virus accessions collected from China and USA are dominant in the ME tree and constantly group with accessions from other countries. Overall, the target SARS-CoV-2 genomes tend to separate into two major clusters (highlighted in yellow dots, **Figure 2**), suggesting these SARS-CoV-2 may have originated from two major spread sources. Of particular interest is the observation of several phylogenetic clades encompassing samples collected from more than one country, which may provide clues to track the spread history of SARS-CoV-2 in these countries. For example, a notable clade (clade a) containing accessions collected from USA, Brazil, Italy, Australia, Sweden, and South Korea was identified. The only Brazil accession (MT126808.1) in this study is found to be clustered with one accession from USA (MT163716.1) with strong support. Whilst the virus accessions from China are prevalent in the ME tree, it is intriguing that no correlated accession from China is found in this clade. An additional clade including accessions collected from China, USA and Finland were found together (clade b). In another notable clade (clade c), 2 of the 3 accessions (LC528232.1 and LC528233.1) collected from the cruise ship in Japan were grouped with several accessions from USA. Two accessions (MT198651.1 and MT198652.1) collected in March 2020 from Spain were grouped (clade f) with one accession collected in January 2020 from China. The additional Spain accession (MT198653.1) was clustered with one from USA (MT192765.1). One India accession (MT012098.1) was found together (clade g) with an accession from Wuhan, China, collected in December 2019. Interestingly, the single Nepal accession (MT072688.1) seems to be closely related (clade d) to several accessions from USA.

### Spike protein of SARS-CoV-2 has undergone a structural rearrangement

The spike glycoprotein is critical for virus infection. Recent study suggested that the S protein in SARS-CoV-2 may have undergone a structural rearrangement^13^. To investigate this hypothesis, two separate phylogenies were developed based on the full-S and RBD sequences, respectively. Human SARS-CoV-2, MERS, and SARS-CoV reference sequences and their close coronavirus homologues identified from various animal hosts were included for the phylogenetic analyses. Overall, the two phylogenies displayed similar clustering patterns, separating into three major clades (**Figure 3)**. SARS-CoV-2 was identified in the same major clade and was clustered most closely with two bat SARS CoVs (highlighted in purple and green colors, **Figure 3**) and the human SARS-CoV (orange color, **Figure 3**). In both phylogenies, SARS-CoV-2 is most closely related to bat_CoV_RaTG13, suggesting SARS-CoV-2 may have originated from bat. However, the evolutionary positions of human SARS-CoV and bat-SL-CoVZ45 were swapped between the full-S and RBD-only phylogenies. In the full-S phylogeny, bat-SL-CoVZ45 is relatively more similar to human SARS-CoV-2, whilst human SARS-CoV is closer to SARS-CoV-2 than bat-SL-CoVZ45. Taken together, these results suggested that the RBD of SARS-CoV-2 is more likely originated from human SARS-CoV, whilst the remaining part of the S protein in SARS-CoV-2 may have originated from bat-SL-CoVZ45, supporting the potential structural rearrangement of S protein in SARS-CoV-2. bat_CoV_RaTG13 is similar to SARS-CoV-2, indicating the proposed structural rearrangement may have occurred in bat first before its transmission to human.

**Figure 3.**
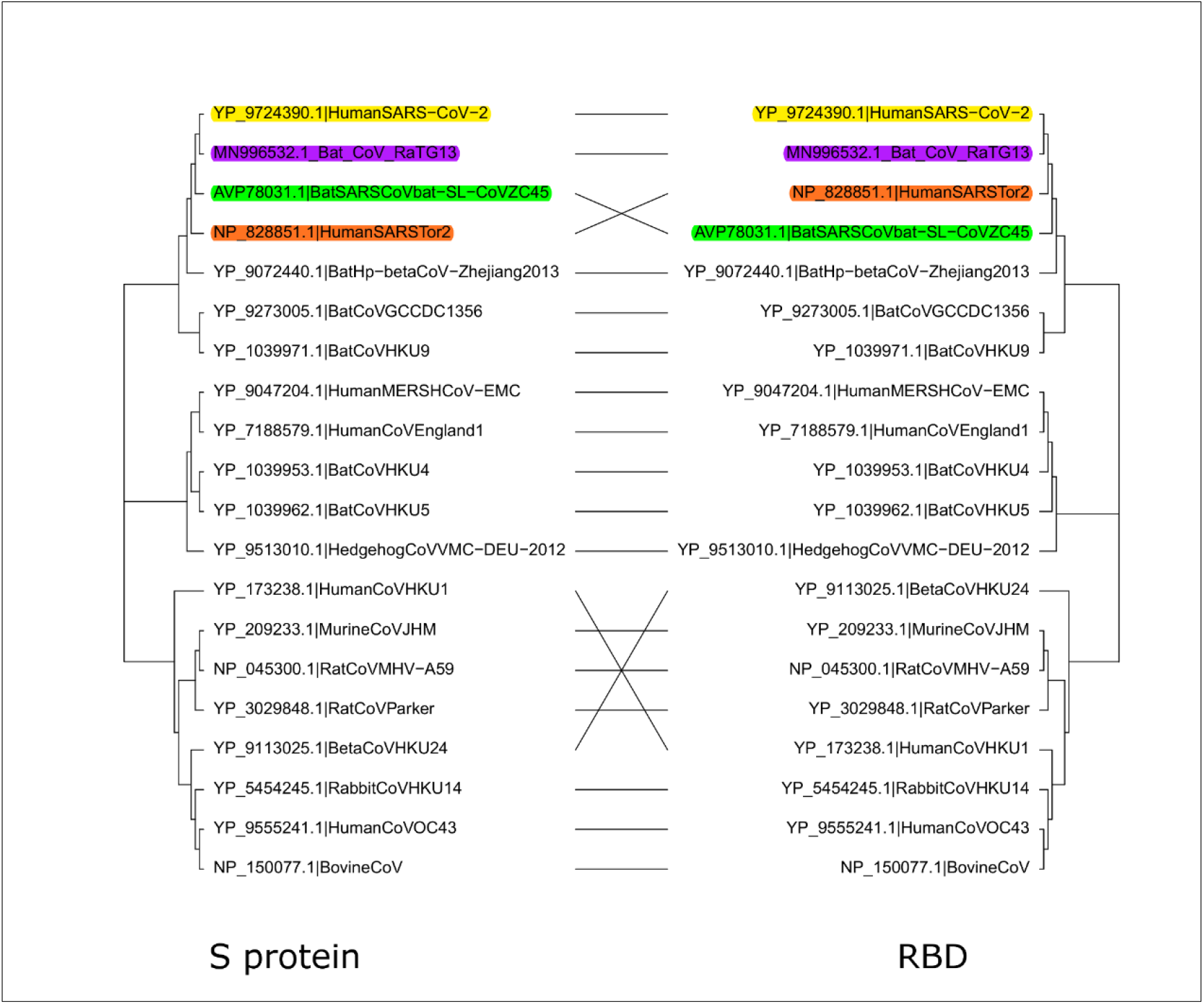
Displays the phylogeny discrepancy of the full-S and RBD sequences. Maximum Likelihood phylogenies based on the full-S protein (left) and RBD (right) sequences of SARS-like CoVs. Taxa names were in the “Accession Id, host organism, sample name” format. Human SARS-CoV-2 and its close relatives were highlighted in different colors.

### A single amino acid mutation in RBD results in reduced receptor binding affinity on human ACE2

The RBD of virus S protein binds to a receptor in host cells, and is responsible for the first step of CoV infection ^3^. Thus, amino acid mutation to RBD may have significant impact on receptor binding and vaccine development. The 3D structure of the spike protein RBD of SARS-CoV-2 (PDB: 6VW1) has recently been determined in complex with human ACE2 receptor ^6^. One of the 12 spike protein mutations identified above (**Figure 1B**) was located in the RBD region (R408I). Further data screening against the latest GISAID and NCBI database (5^th^ Oct) revealed a total of 17 strains from five countries containing the R408I mutation (**Table 1**). Sequence alignment showed that 408R is strictly conserved in SARS-CoV-2, SARS-CoV and bat SARS-like CoV (**Figure 4A**). Based on the determined CoV2_RBD-ACE2 complex structure, 408R is located at the interface between RBD and ACE2, but is positioned relatively far away from ACE2 and thus does not have direct interaction with ACE2 (**Figure 4B**). However, the determined RBD0-ACE2 structure showed that 408R forms a hydrogen bond (3.3 Å in length) with the glycan attached to 90N from ACE2 (**Figure 4C**) ^6^. The hydrogen bond may have contributed to the exceptionally higher ACE2 binding affinity. The arginine residue is also conserved in human SARS-CoV (corresponding to 395R in PDB: 2AJF), but is positioned relatively distant (6.1 Å) from the glycan bound to 90N from ACE2 (**Figure S1**). Interestingly, the 408R-glycan hydrogen bond appears to be disrupted by the R408I mutation in one SARS-CoV-2 accession (GeneBank ID: MT012098.1) (**Figure 4D**), collected from India on 27^th^ Jan 2020. *In silico* calculations indicatethat the R408I mutation increased the binding free energy by 0.93 kcal/mol, predicting a modest decrease in ACE2-binding affinity. In contrast to the electrically charged and highly hydrophilic arginine residue, the mutated isoleucine residue has a highly hydrophobic side chain with no hydrogen-bond potential (**Figure 4E**). In summary, the R408I mutation identified from the SARS-CoV-2 strain in India represents a SARS-CoV-2 mutant with likely lower ACE2 binding affinity.

**Figure 4.**
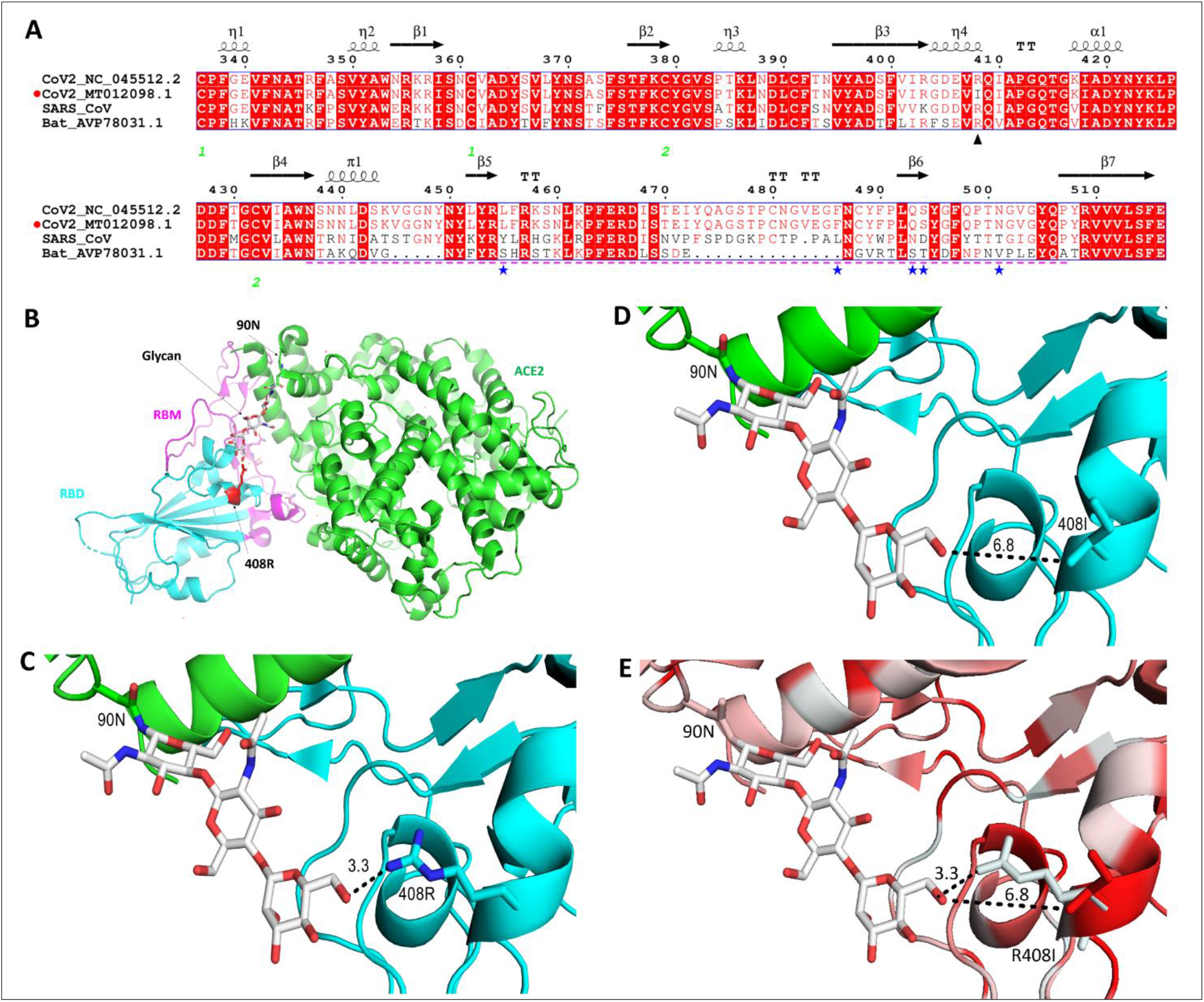
Sequence alignment and protein structural analyses of the mutation in RBD of SARS-CoV-2. **A**) Sequence alignment of RBDs. ▲: R408I mutation; ---: receptor binding motif (RBM). ⋆: RBD-interacting sites. **B**) Overall position of the identified mutation relative to: RBD (cyan), ACE2 (green) with RBM (pink) and Glycan (grey). **C,D**) Display the disrupted hydrogen bond by the R408I mutation. “---”: distance in Å. **E**) Hydrophobic profile changes due to R408I mutation, with red and white colors representing the highest hydrophobicity and the lowest hydrophobicity, respectively. All amino acid number according to the S protein of SARS-CoV-2 (NC_045512.2) and human ACE2, respectively.

## Discussions

Based on the currently available genome sequence data, our results showed that the mutation rate of SARS-CoV-2 is much lower than that for SARS, which caused the 2002-2003 outbreak. Our study is the first to provide a direct gene-based quantitative comparison between SARS-COV-2 and SARS. Among the different genetic regions of SARS-CoV-2 genomes, we found that the spike protein (S) is more conserved that other genetic regions such as ORF1, ORF8, and N, which encode nonstructural polyprotein, virus accessory protein, and the nucleocapsid protein, respectively. A relatively stable spike protein region of SARS-CoV-2 is a good indication for the epidemic control, as less mutation raises the hope of the rapid development of a vaccine and antiviral drugs. Our results are consistent with several recent genetic variance analyses on SARS-CoV-2 ^15,23–25^, which suggested the SARS-CoV-2 genomes are highly homogeneous. Furthermore, based on the latest genomic data for SARS-CoV-2, molecular geneticists monitoring the virus development also suggested that the mutation rate of SARS-CoV-2 maintains at a low level ^17,26,27^. Whilst it is generally safe to say that SARS-CoV-2 tends to mutate at a low rate, as the virus continues to spread rapidly around the world, and more genomic data is accumulated, the evolution and mutation dynamics of SARS-CoV-2 still need to be monitored closely.

One critical aim of our study is to identify the original status of SARS-CoV-2 before its wide transmission across different countries. Due to the short time space of sample collection and a relatively low mutation rate for SARS-CoV-2, we believe that Minimum Evolution phylogeny (a parsimony method) may outperform other phylogenetic methods to achieve this aim. Similarly, Peter et al. ^28^ also adopted a parsimony phylogeny (Maximum Parsimony) to trace the spread history of SARS-CoV-2 in the early stage of the pandemic. Minimum Evolution and Maximum Parsimony are similar phylogeny methods (both using the parsimony sites detected in the sequence alignment) trying to minimize the total number of substitutions in the phylogenetic tree. In our analysis, the earliest few reported SARS-CoV-2 accessions collected from Wuhan China were identified at the center of the phylogenetic tree with the shortest branch. Interestingly, several virus genomes from USA were found to be identical to these putative original versions of the virus from Wuhan. According to public media, the outbreak of SARS-CoV-2 in USA occurred relatively later than other countries. One possible explanation for this observation is that, the spread of SARS-CoV-2 in USA might start much earlier than previously thought or reported. Since a dominant proportion of the samples in this study were collected from China and USA, we observed a significantly higher level of genetic diversity from these two countries. Most SARS-CoV-2 accessions from the other countries can find their closely related sisters from either China or USA. This data bias, on the other hand, may give us an advantage to trace the spread history of SARS-CoV-2 in different countries. This suggestion is reliable because all samples investigated in this study were collected at the early stage of the pandemic, which may avoid the potential data noise caused by recent published genomes of complex spread background. One notable finding in our phylogenetic tree is that, the singleton SARS-CoV-2 accessions collected from Australia, Brazil, South Korea, Italy and Sweden were clustered together with two USA samples but without a Chinese version, suggesting that these infection cases may be somehow related. In addition, one of the three samples collected from the cruise ship stranded in Japan was found to be closely related to a sample collected from Guangzhou, China, whilst the other two were grouped with several cases from USA. Noteworthy, our phylogeny seems to support the presence of two major types of SARS-CoV-2 in the target samples, suggesting the potential existence of two spread sources. Interestingly, this speculation is corroborated by an independent clustering analyses using a different phylogeny method ^23^.

Until now, the origin of SARS-CoV-2, and how it has been transmitted to humans remains largely a mystery. Early genomic data indicated that human SARS-CoV-2 is an enveloped, positive-sense, and single-stranded RNA virus in the subgenus *Sarbecovirus* of the genus *Betacoronavirus* ^13,14^. Evolutionarily, SARS-CoV-2 is most closely related to bat SARS-like CoV (88% genome sequence identity) and human SARS CoV (79%), the latter of which caused a global pandemic in 2003 ^13^. Based on the strong genome sequence identity between SARS-CoV-2 and bat SARS-like COVs, it was initially speculated that SARS-CoV-2 may have originated from bat ^14,29^. However, a more recent study proposed that pangolin may be the most likely reservoir hosts due to the identification of closely related SARS-COVs from this species as well ^30^. Both animals can harbor coronaviruses related to SARS-CoV-2. However, direct evidence of the transmission of SARS-CoV-2 from either bat or pangolin to human is still missing.

Prior to this study, several publications have suggested that SARS-CoV-2 may have originated from the genome recombination of SARS-like CoVs from different animal hosts, as evidenced by the discrepant clustering patterns for the phylogenies using different genetic regions. Lu ^13^ first observed that the RBD of S protein in SARS-CoV-2 is more closely related to human SARS-CoV, whilst the other part of its genome is more similar to bat SARS-CoV. Later Lam ^30^ identified a bat CoV_RaTG13 and several pangolin SARS-CoVs that are consistently closer to SARS-CoV-2 than human SARS-CoV in either full-S protein or RBD. By combining the data from these two studies, our study confirmed the observations reported in both studies, and further determined that the S protein recombination actually happened between human SARS-CoV and a bat SARS-CoV, much earlier before its transmission to human, with the newly identified bat SARS-CoV-RaTG13 as an intermediate.

The RBD of S protein binds to a receptor in host cells and is responsible for the first step of CoV infection. The receptor binding affinity of RBD directly affects virus transmission rate. Thus, it has been the major target for antiviral vaccine and therapeutic development such as SARS ^8^. At the time of first completion in late March 2020, this study was the first to report the identification of the R408I mutation in the RBD of S protein in SARS-CoV-2. Since then, the R408I mutant has attracted research attention from a significant number of researchers. Both computational and experimental studies have been performed to further investigate its molecular characteristics and potential immune effects ^31–38^. In addition, commercial synthesis of the R408I recombinant RBD protein has been offered by serval companies (Acro Biosystems, Creative Dianostics, SinoBiological, and Creative Biolabs) for immuno-binding and diagnostic testing. Noteworthy, Yan et al. ^31^ showed that three of the four RBD neutralizing antibodies tested could not bind the R408I mutant, whereas other mutants displayed strong binding interaction with all the neutralizing antibodies tested. The authors stated that 408R played an essential role for SARS-CoV-2 RBD antibody binding and the R408I could abolish the antibody binding interaction ^31^. In addition, Zhe et al. ^39^ also suggested that R408I constitute the RBD epitope residues. These observations contrast an early stage study ^17^ which did not notice the R408I mutation and predicted that a single vaccine may be sufficient for all circulating SARS-CoV-2 variant. Based on the determined RBD-hACE2 protein structure (PDB: 6VW1) ^6^, we found that 408R residue can establish an indirect receptor interaction via a glycan attached to human ACE2. This residue was found to be conserved in SARS and MERS as well. Interestingly, the arginine residue (corresponding to 395R in SARS) has also been shown to be involved in receptor interaction in SARS. In this study, we were the first to show that the R408I mutation in SARS-CoV-2 is likely to cause a reduced binding affinity to human ACE2 receptor. Our result has been corroborated in several independent studies later on ^32–34^. Although the S protein gene seems to be more conserved than the other protein-encoding genes in the SARS-CoV-2 genome, our study provides direct evidence that a mutated version of SARS-CoV-2 S protein with varied transmission rate may have already emerged. Furthermore, we confirmed that, as of 5^th^ Oct 2020, a total of 17 SARS-CoV-2 strains containing the R408I mutation were present in the GISAID and NCBI databases, with the latest R408I mutants collected on 10^th^ Sep 2020 and 1^st^ Sep 2020 from Switzerland and England, respectively. These results suggest that the R408I mutant may spread across to different countries since its first emergence from India and is still actively spreading in different regions. Benson et al.^40^ recently reported that R408I accounts for ~2% of infection cases in Africa. We believe that the number of identified R408I mutants are still underestimated, given the limited sequencing capability in respective countries. Based on the close relationship of SARS-CoV-2 to SARS, current vaccine and drug development for SARS-CoV-2 has also focused on the S protein and its human binding receptor ACE2 ^7,41^. Considering the significantly varied antibody binding profile for R408I, we propose that this mutant still requires significant attention from doctors and scientists around the world during the development of SARS-CoV-2 therapeutic solutions. One suggestion for the next step of therapeutic development is to focus on the identification of potential human ACE2 receptor blocker, as suggested in a recent commentary ^7^. This approach will avoid the above-mentioned challenge faced by vaccine development.

## Acknowledgement

The authors would like to thanks the relevant research community for making the genomic data available to the public. We thanks Dr Yinchuan Zhang from Maternal and Child Health Hospital, Dingxi, Gansu, China, and Mrs Hong Ma from Centre of Disease Control (CDC), Dingxi, Gansu, China for providing critical comments on the manuscript.

## Author Contribution

WLW, CL and YJ conceived the study. YJ, GS, SN, JBB, YZ, KSH, HYH, WSH, CHY performed data analyses. YJ, GS&SN wrote the manuscript. All authors have read the manuscript.

## Supplementary figure

**Figure S1.**
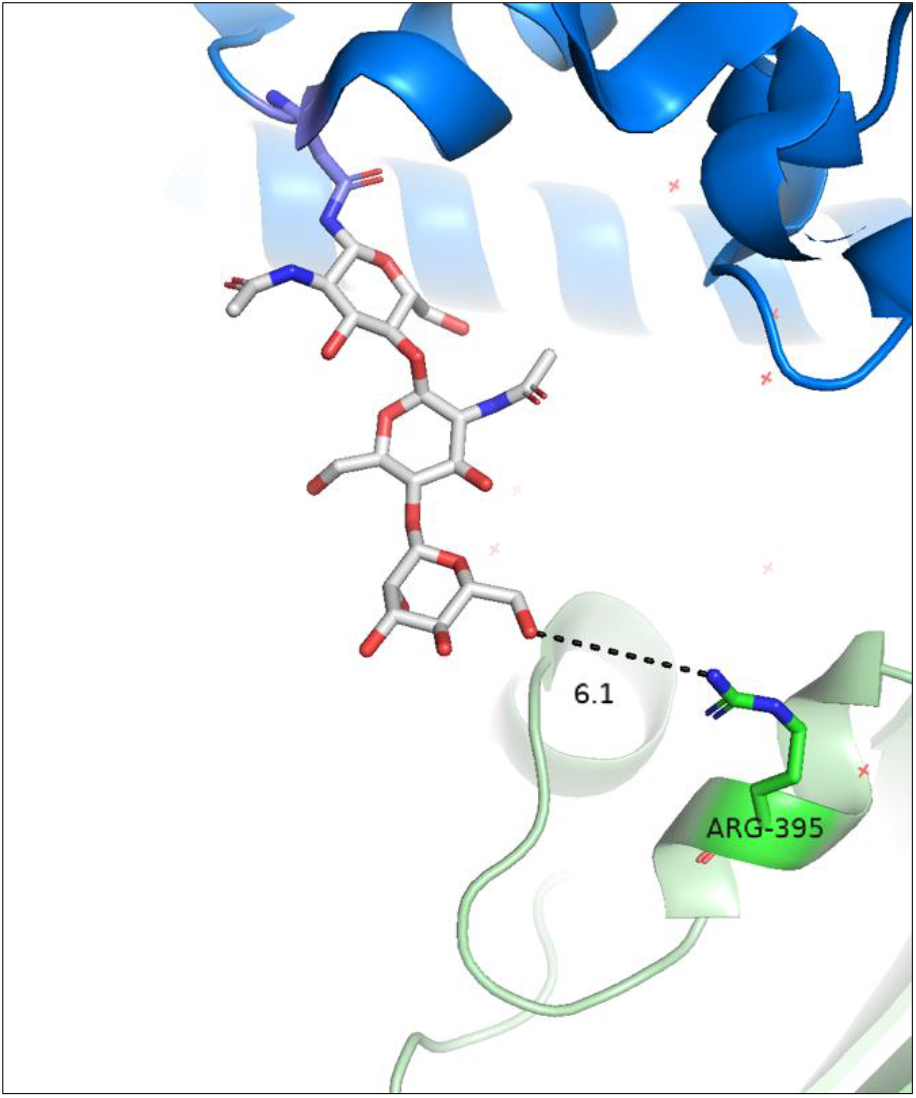
Displays the position of 395R in human SARS-CoV (PDB: 2AJF). Dash line indicates the measured distance in Å.

## Data availability

The Genebank ID list of 106 SARS-CoV-2 and 39 SARS genomes used in this study is available at https://figshare.com/s/3d3c24ef05084b534b4c

